## Supplementary Data 1: List of software packages and dependencies for "OmnibusX: A unified platform for accessible multi-omics analysis"

|  | **Package/Library** | **Repository URL** |
| --- | --- | --- |
| 1 | Python | <https://github.com/python/cpython> |
| 2 | FastAPI | <https://github.com/tiangolo/fastapi> |
| 3 | Uvicorn | <https://github.com/encode/uvicorn> |
| 4 | Gunicorn | <https://github.com/benoitc/gunicorn> |
| 5 | python-jose | <https://github.com/mpdavis/python-jose> |
| 6 | Requests | <https://github.com/psf/requests> |
| 7 | msgpack | <https://github.com/msgpack/msgpack-python> |
| 8 | SQLModel | <https://github.com/tiangolo/sqlmodel> |
| 9 | psutil | <https://github.com/giampaolo/psutil> |
| 10 | NumPy | <https://github.com/numpy/numpy> |
| 11 | Scanpy | <https://github.com/scverse/scanpy> |
| 12 | SciPy | <https://github.com/scipy/scipy> |
| 13 | h5py | <https://github.com/h5py/h5py> |
| 14 | Pandas | <https://github.com/pandas-dev/pandas> |
| 15 | PyArrow | <https://github.com/apache/arrow> |
| 16 | DoubletDetection | <https://github.com/JonathanShor/DoubletDetection> |
| 17 | pynndescent | <https://github.com/lmcinnes/pynndescent> |
| 18 | cryptography | <https://github.com/pyca/cryptography> |
| 19 | statsmodels | <https://github.com/statsmodels/statsmodels> |
| 20 | harmonypy | <https://github.com/slowkow/harmonypy> |
| 21 | fastcluster | <https://github.com/dmuellner/fastcluster> |
| 22 | scikit-image | <https://github.com/scikit-image/scikit-image> |
| 23 | rdata | <https://github.com/has2k1/rdata> |
| 24 | tifffile | <https://github.com/cgohlke/tifffile> |
| 25 | imagecodecs | <https://github.com/cgohlke/imagecodecs> |
| 26 | opencv-python-headless | <https://github.com/opencv/opencv-python> |
| 27 | openslide-python | <https://github.com/openslide/openslide-python> |
| 28 | pyvips | <https://github.com/libvips/pyvips> |
| 29 | pydeseq2 | <https://github.com/owkin/PyDESeq2> |
| 30 | intervaltree | <https://github.com/chaimleib/intervaltree> |
| 31 | xmltodict | <https://github.com/martinblech/xmltodict> |
| 32 | openpyxl | <https://github.com/chronossc/openpyxl> |
| 33 | aiofiles | <https://github.com/Tinche/aiofiles> |
| 34 | python-multipart | <https://github.com/andrew-d/python-multipart> |
| 35 | requests-toolbelt | <https://github.com/requests/toolbelt> |
| 36 | gseapy | <https://github.com/zqfang/GSEApy> |
| 37 | weblogo | <https://github.com/WebLogo/weblogo> |
| 38 | mellon | <https://github.com/pythons/mellon> |
| 39 | palantir | <https://github.com/dpeerlab/Palantir> |
| 40 | kdepy | <https://github.com/tommyod/KDEpy> |
| 41 | ReactJS | <https://github.com/facebook/react> |
| 42 | Electron | [https://www.electronjs.org](https://www.electronjs.org/) |
| 43 | ThreeJS | <https://threejs.org/> |
| 44 | OpenSeaDragon | [https://openseadragon.github.io](https://openseadragon.github.io/) |
| 45 | ECharts | <https://echarts.apache.org/en/index.html> |
| 46 | D3JS | [https://d3js.org](https://d3js.org/) |
